# The increased production of monocytes with immunosuppressive transcriptional profiles predicts the development of SIV encephalitis

**DOI:** 10.64898/2026.10.14.759063

**Authors:** Hai Duc Nguyen, Andrew K. Ding, Cinar Efe Sumer, Patrick Autissier, Melanie N. Cash, Marco Salemi, Kenneth C. Williams, Woong-Ki Kim

**Affiliations:** Division of Microbiology, Tulane National Biomedical Research Center, Tulane University, Covington, Louisiana, USA; Biology Department, Boston College, Chestnut Hill, Massachusetts, USA; Department of Epidemiology, University of Florida College of Medicine, Gainesville, Florida, USA; Department of Microbiology & Immunology, Tulane University School of Medicine, New Orleans, Louisiana, USA

**Author notes:** **Corresponding authors:** Kenneth C. Wiliams, Biology Department, Boston College, 140 Commonwealth Avenue, Chestnut Hill, MA 02467, USA Woong-Ki Kim, Tulane National Biomedical Research Center, 18703 Three Rivers Road, Covington, LA 70433, USA. These authors contributed equally to this work (Hai Duc Nguyen, Andrew K. Ding).

**Keywords:** BrdU, HAND, HIV, monocytes, neuroinflammation, PARP2

## Abstract

HIV-associated neurocognitive disorders (HAND) remain common despite antiretroviral therapy. We previously demonstrated that increased percentages of newly generated BrdU-labeled (BrdU⁺) monocytes predict AIDS progression and SIV encephalitis (SIVE) as early as 8 and consistently by 21 days post-infection (DPI) in SIV-infected macaques, but their molecular identity remained unknown. We performed Bulk RNA sequencing of FACS-sorted BrdU⁺ and BrdU⁻ monocytes from uninfected animals and SIV-infected macaques at 21 DPI and necropsies, including SIVE or not (SIVnoE). BrdU⁺ monocytes underwent progressive transcriptional remodeling during SIV infection. Early SIVE-predictive monocytes exhibited suppressed interferon, TNF-α/NF-κB, antigen-presentation, and antimicrobial pathways, with increased LILRB4 and PARP2 expression. At terminal disease, SIVE BrdU⁺ monocytes displayed an immunosuppressive profile, reduced HLA-DR expression, and increased structural variants compared with SIVnoE monocytes. Together, newly generated circulating monocytes in animals progressing to SIVE are an immature and immunosuppressive population that emerges before neuropathology, implicating dysregulated monocytopoiesis and pathogenic monocytes in HAND.

## Introduction

In the era of antiretroviral therapy (ART), HIV-associated comorbidities have become the major causes of morbidity and mortality from HIV infection (Trickey et al., 2024). HIV-associated neurocognitive disorders (HAND) and cardiovascular disease (CVD) are the most prevalent HIV-associated comorbidities and affect an estimated 50% of people living with HIV (PLWH) (Wang et al., 2020). Soluble CD163 (sCD163), a biomarker produced exclusively by monocytes and macrophages, is consistently demonstrated to be a biomarker of HAND and CVD compared to general inflammatory biomarkers and classical markers of HIV progression, underscoring the importance of the expansion of activated monocytes/macrophage from bone marrow (BM) and their accumulation in target tissues (Burdo et al., 2013b; Ding et al., 2024; Williams et al., 2020).

Recent reports with regard to hematopoiesis point to monocytes produced in response to infection and injury having distinct developmental origins, functions, and fates, which can both expand immune activation but also potentially dampen immune response (Swann et al., 2024). Using intravenous (i.v.) BrdU administration to label recently produced monocytes in a macaque model of HIV infection, we have shown that an increased percentage and absolute number of BrdU-labeled (BrdU+) monocytes correlate with the rate of AIDS development, the prevalence of SIV encephalitis (SIVE), and the histopathologic severity of macrophage inflammation.

Importantly, such BrdU numbers can differentiate these changes as early as 8 days and consistently 21 days post-infection (DPI) (Burdo et al., 2010). We have more recently shown that monocyte turnover, demonstrated by %BrdU+ monocytes, is a better predictor of AIDS and SIVE, and cardiovascular pathogenesis compared to general inflammatory biomarkers (galectin-3, galectin-9, IL-18), and is linearly correlated with sCD163 levels in serum of infected monkeys (Burdo et al., 2010; Ding et al., 2024). However, the identity and molecular signals and functions of SIVE-predictive BrdU+ monocytes have not been studied. Here, we sought to define the transcriptional identity of BrdU+ monocytes in SIV-infected monkeys with AIDS progressing to SIVE, compared to SIV-infected animals with AIDS and no SIVE (SIVnoE) and non-infected controls. We hypothesized that BrdU+ monocytes in SIVE progressors and SIVE animals are transcriptionally distinct from their SIVnoE counterparts.

We assessed the transcriptomes of SIVE-predictive BrdU+ monocytes by performing bulk RNA sequencing (RNA-seq) on FACS-sorted BrdU+ and BrdU-monocytes collected during acute infection (21 DPI) and with terminal AIDS from SIVE animals, SIVnoE animals, and non-infected controls. In SIVE-progressors, we observed paradoxical downregulation of inflammatory pathways concurrent with peak viral replication and upregulation of the monocyte activation inhibitor LILRB4. Terminally, BrdU+ monocytes from SIVE, but not SIVnoE animals, have upregulated genes associated with immunosuppressive monocyte phenotypes relative to non-infected controls. SIVE BrdU+ monocytes also have upregulation of the DNA repair enzyme, PARP2, and a greater burden of structural variants relative to their SIVnoE counterparts, raising the possibility that aberrant monocytopoiesis is related to immunosuppressive transcriptional profiles in SIVE-predictive and SIVE BrdU+ monocytes.

These findings provide the first transcriptional evidence for the role of immunosuppressive monocytes in AIDS and SIVE pathogenesis, potentially highlighting a role for immunosuppressive monocytes in HAND that may function to downregulate or shut off macrophage innate immune responses once acquired immune responses are dysregulated from HIV infection.

## Results

In this study, we utilized n=7 SIVmac251-infected, CD8-depleted RMs and n=6 non-infected, CD8-depleted controls (**Table S1**). CD8-lymphocyte depletion was greater than 21 days in all animals. Of the infected animals, n=3 developed SIVE and n=4 did not develop SIVE (SIVnoE). For transcriptome studies, BrdU+ monocyte numbers and percentages were assessed in whole blood by analytical flow cytometry, from the animals at 24 h post i.v BrdU inoculation.

Additionally, BrdU+ and BrdU-monocytes were collected from PBMCs and recovered by flow cytometry sorting at 21 DPI and at necropsy and used for subsequent bulk RNA-seq (**Fig. S1, S2, S3, S4, and S5**). Sorting purity for all samples exceeded 99% (**Fig. S6**). Longitudinal BrdU labeling revealed increased monocyte turnover during early SIV infection, with the frequency of BrdU⁺ monocytes at 21 DPI distinguishing animals that progressed to SIVE from those that did not (estimated difference = 18.225, p = 0.031), whereas non-infected controls remained at baseline levels of approximately 5% or less (**Fig. 1A-B**). The 21 DPI time point was chosen because we have previously shown that the %BrdU+ monocytes predict SIVE progression as early as 21 DPI, and the necropsy time point was chosen because SIVE histopathology develops terminally and the percentage of BrdU+ cells correlates with the rapidity of AIDS development and severity of CNS histopathology with SIVE. **Fig. 2A–D** illustrate the transcriptional changes in BrdU⁺ monocytes across disease stages. Compared with uninfected controls, BrdU⁺ monocytes from SIV-infected rhesus macaques exhibited marked transcriptional alterations. A core set of 29 genes was significantly downregulated in BrdU⁺ monocytes across all three conditions (21 DPI, SIVnoE, and SIVE) (**Fig. S7A,B** and **Table S2**). Enrichment analysis showed that these genes were involved in metabolism, inflammation, immune response, HIV infection and HAND pathology, immunology, and gene regulation (**Fig. S7C**). These unique sets point to stage- and pathology-specific transcriptional adaptations in proliferating monocytes. Notably, BrdU⁺ monocytes from animals with SIVE displayed a unique 10-gene signature that distinguished them from both SIVnoE animals and uninfected controls, indicating disease-specific transcriptional reprogramming. These genes are associated with biological processes relevant to SIVE, including immune cell migration (FMNL2, SPN), inflammatory signaling (TRIM47), and regulation of cell proliferation (CDKN1C) (**Fig. 2E**).

**Fig. 1.**
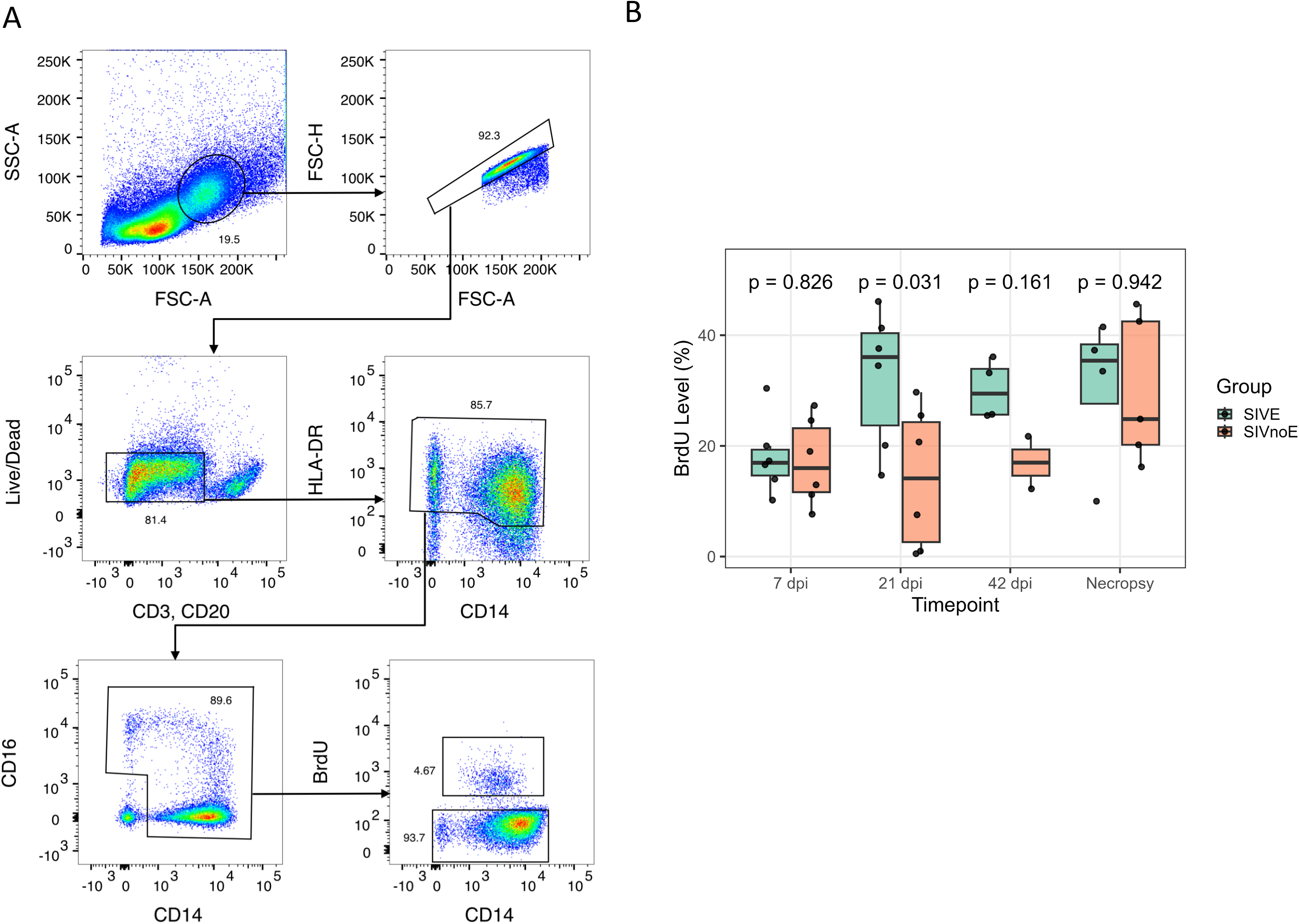
Longitudinal sampling of BrdU+ monocytes. BrdU+ monocytes were measured throughout infection in SIV-infected, CD8+ T-Cell-depleted rhesus macaques with matched timepoints in non-infected, CD8+ T-Cell-depleted controls. **A)** At 21 days post-infection and at necropsy, BrdU+ and BrdU-monocytes were sorted from PBMCs. Monocytes were first identified by forward and side scatter profiles (FSC, SSC, respectively), doublets were excluded by an FSC-area vs. height plot, and dead cells removed using a cell viability dye. Next, monocytes were identified based on expression of HLA-DR, CD14, and CD16, then sorted into BrdU+ and BrdU-fractions. **(B)** Longitudinal BrdU⁺ monocyte frequencies in blood of SIV-infected animals progressing to SIVE (SIVE) or not developing SIVE (SIVnoE) and non-infected controls. BrdU⁺ monocytes were measured at 7, 21, and 42 DPI and at necropsy, with group differences evaluated at each timepoint using two-way ANOVA followed by Tukey-adjusted pairwise comparisons. Data are shown as individual animals with boxplots indicating the median and interquartile range.

**Fig. 2.**
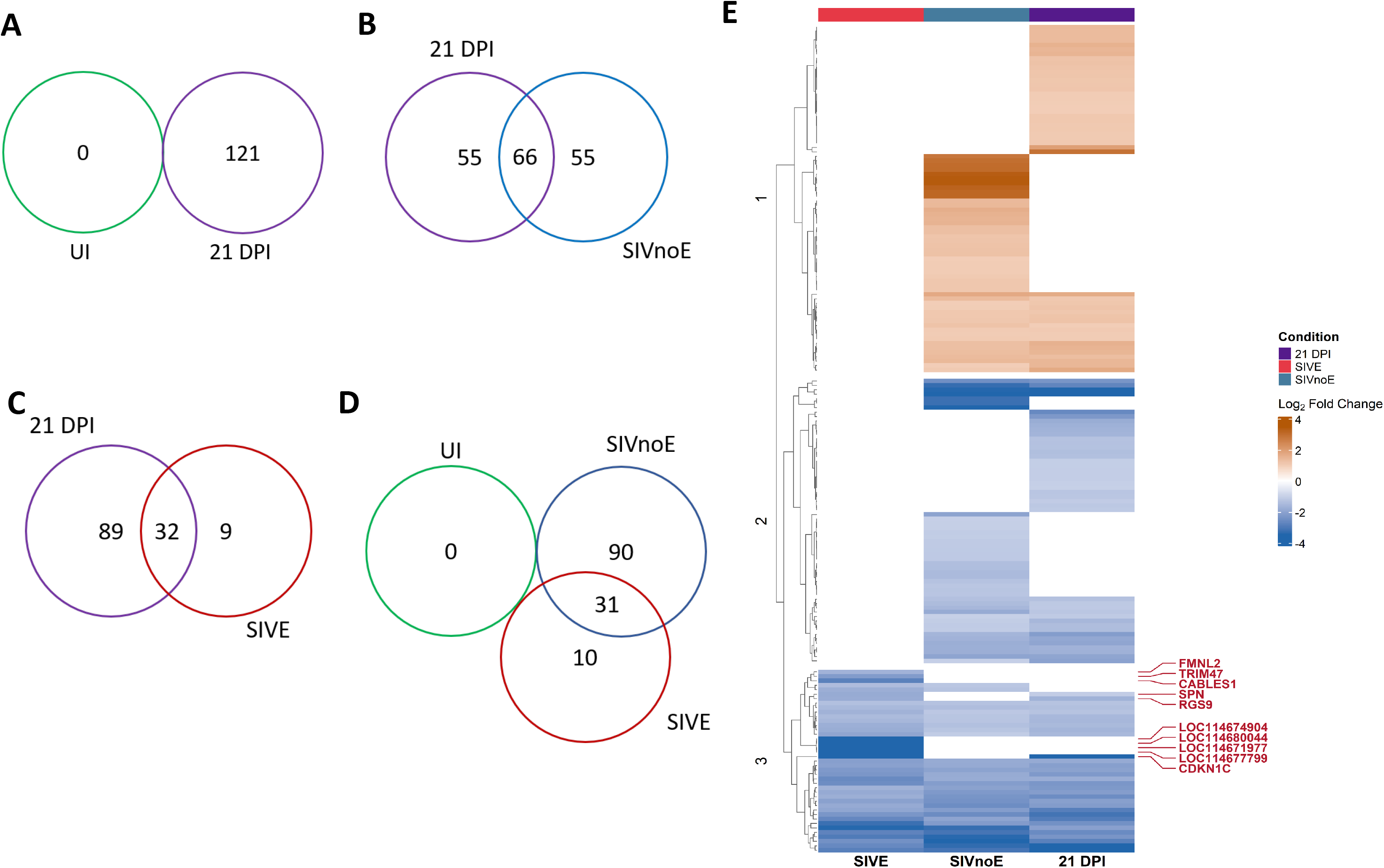
Transcriptional changes across disease stages. Venn diagrams showing differentially expressed genes (DEGs) identified in comparisons between (A) uninfected controls and 21 DPI, (B) 21 DPI and SIVnoE, (C) 21 DPI and SIVE, and (D) across uninfected controls, 21 DPI, and SIVnoE. (E) Heatmap showing distinct and shared DEGs across disease stages, highlighting the 10 genes uniquely associated with SIVE.

**Fig. 3.**
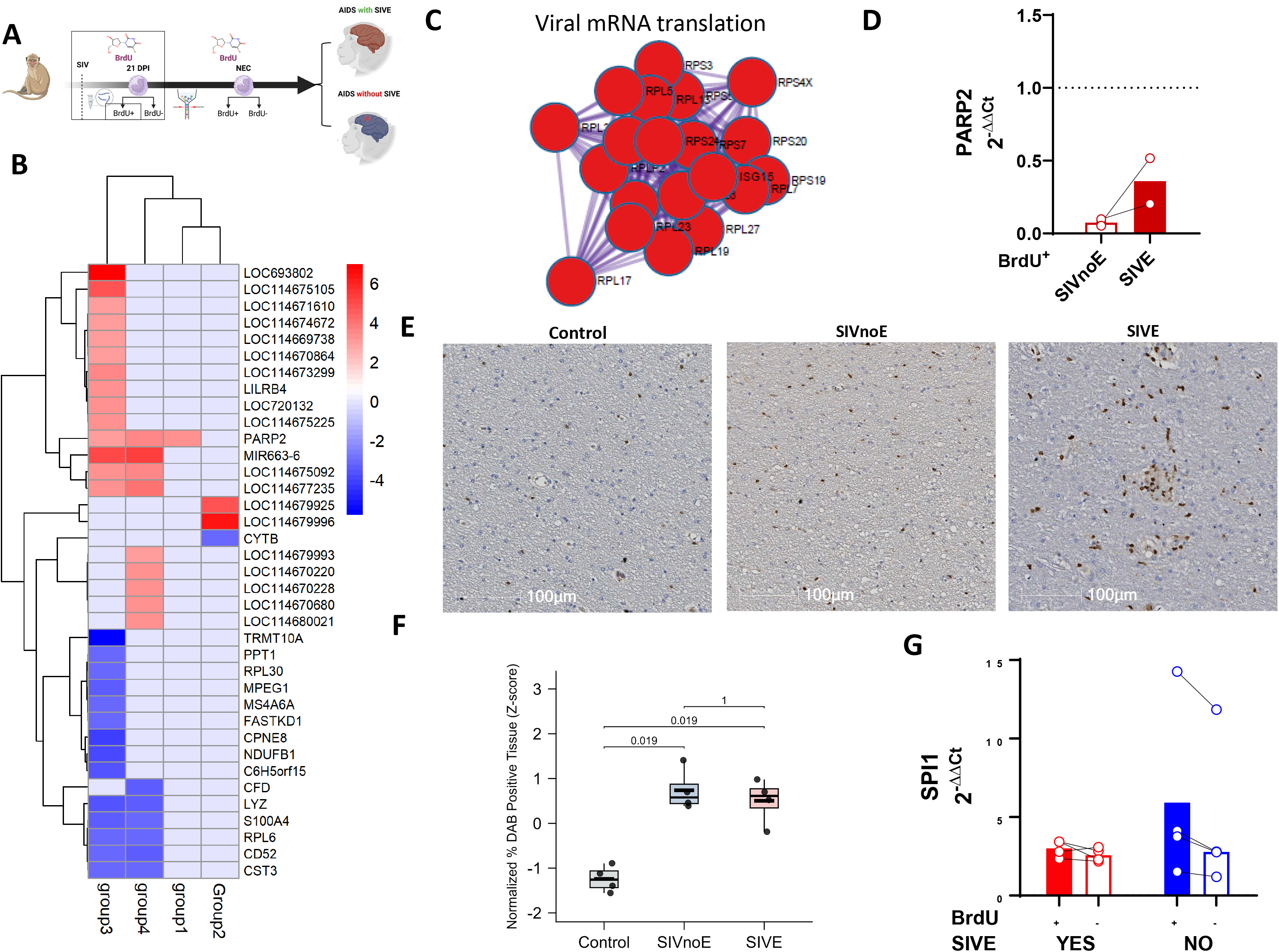
Early transcriptional signatures in BrdU⁺ monocytes are associated with progression to SIV-associated encephalitis. (**A**) Experimental groups: Group 1 (SIVE progressors) and Groups 2–4 (SIVnoE non-progressors) were administered BrdU at 21 days post-infection (DPI). (**B**) Heatmap of selected differentially expressed genes (DEGs) across groups. (**C**) Pathway enrichment analysis (log₁₀ *P* values) showing suppression of pathways including viral mRNA translation (log₁₀ *P* = −47.7), identified using Metascape with MCODE (Molecular Complex Detection) to detect densely connected subnetworks within protein–protein interaction networks. (**D**) qPCR analysis of *PARP2* expression in BrdU⁺ monocytes from SIVnoE and SIVE animals. (**E**) Immunohistochemistry (IHC) of SPI1/PU.1 in brain tissue from control, SIVnoE, and SIVE macaques. (**F**) Quantification of SPI1/PU.1 expression across groups using one-way ANOVA followed by Tukey’s multiple comparisons test. (**G**) qPCR analysis of *SPI1* expression in BrdU⁺ versus BrdU⁻ monocytes in SIVE and SIVnoE animals.

**Fig. 4.**
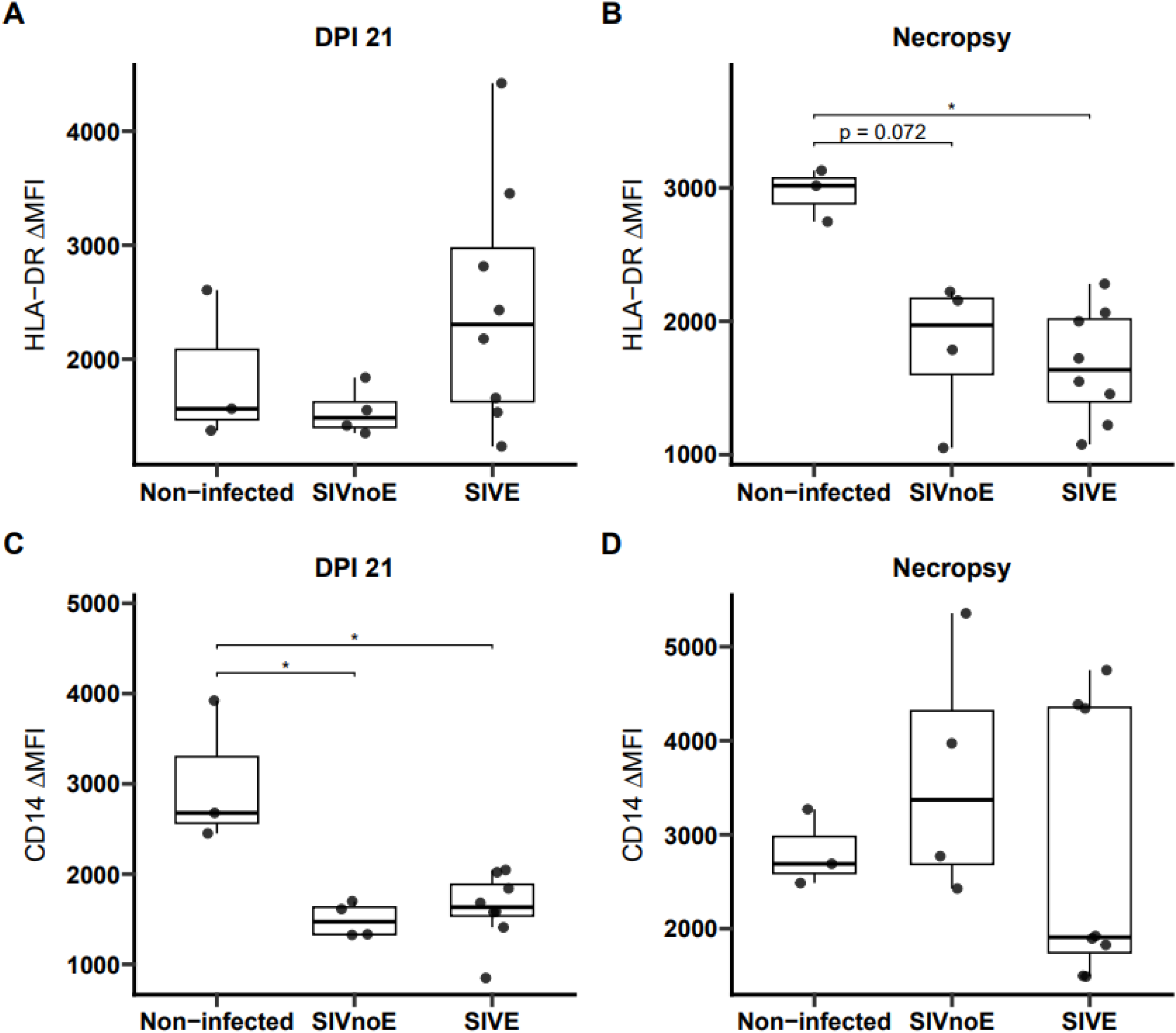
Surface phenotype of newly generated (BrdU⁺) monocytes during SIV infection. **(A)** HLA-DR expression on BrdU⁺ monocytes at 21 days post-infection (DPI) in non-infected controls, SIV-infected animals that later developed SIVE (SIVE progressors), and SIV-infected animals that did not develop SIVE (SIVnoE). **(B)** HLA-DR expression on BrdU⁺ monocytes at necropsy. **(C)** CD14 expression on BrdU⁺ monocytes at 21 DPI. **(D)** CD14 expression on BrdU⁺ monocytes at necropsy. Data are presented as individual animals with mean ± SEM. Statistical significance was determined using one-way ANOVA with appropriate multiple-comparison testing. *P* < 0.05; ns, not significant.

To begin to characterize the transcriptomes of BrdU+ monocytes in SIV infection, we compared the transcriptomes of BrdU+ to BrdU-monocytes within SIVE and SIVnoE animals at necropsy (**Fig. S1 and S2**). In animals with SIVE, BrdU+ monocytes downregulated 41 transcripts relative to BrdU-monocytes, including CX3CR1, CFD, and MS4A7 (**Fig. S1D** and **Table S3**). GO analysis showed that downregulated genes were enriched for processes including inflammatory response, mononuclear cell proliferation, and leukocyte migration (**Fig. S1F**). In SIVnoE animals, BrdU+ monocytes downregulated 72 transcripts and upregulated 49 transcripts (**Fig. S2D** and **Table S4**). We observed overlaps with SIVE animals in downregulated transcripts, including CX3CR1, CFD, and MS4A7. In contrast to SIVE animals, BrdU+ monocytes in SIVnoE animals upregulated a number of pro-inflammatory mediators relative to BrdU-monocytes, including CXCL8 and S100A8 (**Fig. S2D**).

We next compared the transcriptomes of BrdU+ monocytes at necropsy between SIVE, SIVnoE, and non-infected control animals (**Fig. S5A**). We identified 8 differentially expressed genes (DEGs) in the SIVE vs. non-infected animals and 144 DEGs in the SIVnoE vs. non-infected comparison (**Fig. S5B** and **Table S5**). When compared to non-infected BrdU+ monocytes at necropsy, SIVE BrdU+ monocytes upregulated genes associated with immunosuppressive monocyte phenotypes, including IDO1, FAM20A, and CD13, and downregulated CD56 (**Fig. S5C**). When compared to SIVnoE BrdU+ monocytes at necropsy, SIVE BrdU+ monocytes upregulated SPC25, LRCH2, BBOX1, ZMAT4, MSX1, and TMEM116 (**Fig. S5C**). We observed that BrdU+ monocytes from SIVE and SIVnoE animals only shared one DEG (*LOC718510*) relative to non-infected animals, and we identified eight DEGs between SIVE and SIVnoE animals (**Fig. S5B**), consistent with the notion that BrdU+ monocytes in SIVE animals are transcriptionally distinct from BrdU+ monocytes in SIVnoE animals.

We have previously shown that an increased %BrdU+ of monocytes is a sensitive and reliable predictor of the rate of progression to AIDS and SIVE histopathologic severity (Hasegawa et al., 2009). Principal component analysis (PCA) demonstrated incomplete separation between groups, except for 2 samples (**Fig. S3B**). A heatmap revealing the top 50 genes across conditions is presented in **Figure S3C**. A total of 121 DEGs (74 downregulated and 47 upregulated) were observed. As observed in chronic SIVE and SIVnoE animals, the majority of DEGs were downregulated in BrdU⁺ relative to BrdU⁻ monocytes (**Table S6**). The most downregulated transcripts included FCGR3 (logFC = -3.59, adj.P.Val = 2.92e-06), CDKN1C (logFC = -3.49, adj.P.Val = 0.0017), HES4 (logFC = -3.62, adj.P.Val = 0.0077), CX3CR1 (logFC = -2.45, adj.P.Val = 0.0018), CFD (logFC = -2.32, adj.P.Val = 0.0048), and ITGAD (logFC = - 2.15, adj.P.Val = 0.00074). Additional downregulated genes of interest included LPL, MS4A7, PLD4, NGFR, ITGAL (logFC = -1.30, adj.P.Val = 0.0017), and the chemokine CXCL10 (logFC = -1.97, adj.P.Val = 0.0129). Furthermore, BrdU⁺ monocytes underwent extensive transcriptional reprogramming between early (21 DPI) and terminal SIV infection, with widespread downregulation of genes and distinct separation of the two stages by PCA. Terminal-stage monocytes exhibited enrichment of neuronal, synaptic, and cell adhesion pathways and were characterized by a regulatory network centered on hub genes such as CAV1, TGFB1, and RHOA, together with activation of neuronal-associated transcription factors, indicating profound functional remodeling during disease progression (**Fig. S4** and **Table S7**).

To identify transcriptional features of BrdU+ monocytes that predict SIVE development, we compared the transcriptomes of BrdU+ monocytes from animals that progressed to SIVE to animals without SIVE (**Fig. 3A**). SIVE-predictive BrdU+ monocytes downregulated lysosomal and antimicrobial genes (LYZ, S100A4, MPEG1), interferon signaling (ISG15, OAS1/2, IFI6, IFIT3), and cytoskeletal regulators (ACTG1, CFL1, PFN1) (**Fig. 3B** and **Table S8**). These downregulated genes were enriched for pathways including interferon α/β signaling (R-HSA-909733, Log10(P)=-5.96), and TNF-α/NF-κB pathways (CORUM:5266, Log10(P)=-7.31).

Protein–protein interaction network analysis further identified significant enrichment of proteins involved in viral mRNA translation (R-HSA-192823, log10(P) = −47.7) (**Fig. 3C**). SIVE-predictive BrdU+ monocytes upregulated the monocyte activation inhibitor LILRB4, the DNA repair enzyme PARP2, along with ADAM8, ADAM28, and MIR663-6 (**Fig. 3B**). PARP2 expression was also higher in BrdU⁻ compared to BrdU⁺ monocytes, as confirmed by PCR (**Fig. 3D**).

To determine whether the transcriptional profile of SIVE-predictive and SIVE BrdU+ monocytes was accompanied by an immature surface phenotype, we assessed HLA-DR and CD14 surface expression. For these analyses, we used data from the animals used for RNA-seq and in historical cohorts for which flow cytometry data was available. At necropsy, HLA-DR expression in SIVE BrdU+ monocytes was significantly lower than non-infected controls (p = 0.0279) and trended lower for SIVnoE BrdU+ monocytes (p = 0.072) versus non-infected controls. HLA-DR expression did not distinguish between SIVE and SIVnoE BrdU+ monocytes (p = 0.494). At 21DPI, HLA-DR expression was similar between BrdU+ monocytes from SIVE progressors, animals that did not progress to SIVE, and non-infected controls. CD14 expression was similar in BrdU+ monocytes across groups at necropsy, but significantly lower in BrdU+ monocytes from SIVE progressors (p = 0.0347) and non-progressors (p = 0.0312) relative to controls (**Fig. 4A-D**).

In earlier comparisons, we observed that the monocyte/macrophage maturation factor PU.1 was a central regulator of differentially expressed genes across comparisons (**Fig. S1G** and **S2G**), raising the possibility that dysregulated monopoiesis contributed to transcriptional changes observed in BrdU+ monocytes. Transcription factor analysis identified SPI1 as a recurrent regulator across DEG comparisons and prior analyses. In brain tissues from control, SIVnoE, and SIVE macaques, SPI1 expression was significantly higher in the SIVnoE, and SIVE groups compared to controls (ANOVA, p = 0.019), as measured by immunohistochemistry (**Fig. 2E–F**). By qPCR, we observed that PU.1 mRNA levels were increased in BrdU-monocytes relative to BrdU+ monocytes in SIVnoE animals but were similarly low in BrdU+ and BrdU-monocytes of SIVE animals (**Fig. 2G**). Structural variant (SV) analyses identified a greater number of SVs in terminal SIVE BrdU+ monocytes compared to those at 21 DPI or to SIVnoE BrdU+ monocytes both at 21 DPI and necropsy. The number of SVs in terminal SIVE BrdU+ monocytes was approximately 3-fold higher than in non-infected animals, which remained consistently low from the 21 DPI and the terminal timepoints (**Fig. 5A–C**). We identified eight SVs that were exclusive to SIVE BrdU+ monocytes – MACROD2, LRP1B, ROBO2, WWOX, PTPRN2, CSMD1, CNTNAP2, and RBFOX1 (**Fig. 5D**) - a relative increase in RYR2 and SMYD3 SV counts in terminal SIVE BrdU+ monocytes compared to their SIVnoE counterparts (**Fig. 5E** and **Table S9**). Among these, SMYD3 emerged as the most consistently altered candidate (**Fig. S8**). Network analysis of SV-harboring DEGs identified hub genes including *BCL2, GSK3B, APP, ESR1*, and *EGFR*, which are associated with cell survival, immune regulation, and neurodegeneration (**Fig. 5F**). Functional enrichment further revealed significant association with neuronal development and CNS injury pathways (**Fig. 5G**). These results are consistent with a model where prolonged monopoiesis produces structural variation that results in inflammatory defects in SIVE BrdU+ monocytes.

**Fig. 5.**
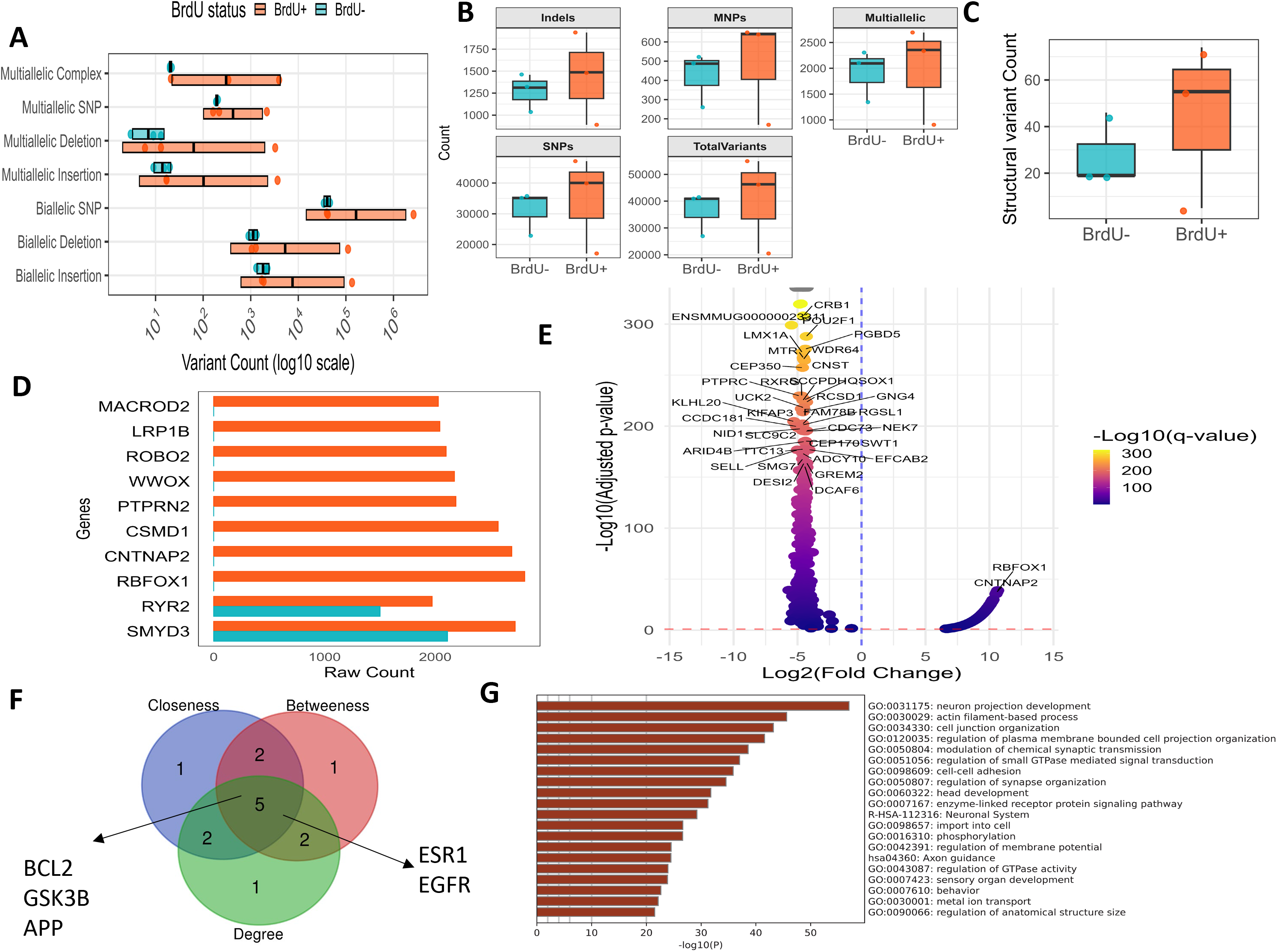
Increased structural variant burden in proliferating monocytes in SIV-associated encephalitis. (**A–C**) Number of structural variants (SVs) detected by DeepVariant (<50 bp; **A**), and FreeBayes (<50 bp; **B**), and Manta (>50 bp; **C**), in BrdU⁺ versus BrdU⁻ monocytes across SIVE, SIVnoE, and 21 DPI. (**D**) Top 10 genes harboring SVs based on SnpEff annotation. (**E**) Genes with significantly different mutation burdens between BrdU⁺ and BrdU⁻ cells, assessed using Fisher’s exact test with Benjamini–Hochberg false discovery rate (FDR) correction. (**F)** Network analysis of SV-harboring DEGs identifying hub genes. (**G**) Functional enrichment analysis of SV-associated genes using Metascape (values represent −log₁₀ *P*).

## Discussion

In this study, we defined the transcriptomic landscape of newly generated (BrdU⁺) monocytes during SIV infection and identified molecular features associated with disease progression and SIVE. We found that BrdU⁺ monocytes undergo progressive transcriptional reprogramming during infection, acquire a distinct SIVE-predictive gene signature before the onset of neuropathology, display an immature surface phenotype characterized by reduced HLA-DR expression, and accumulate structural variants during terminal disease. Together, these findings demonstrate that newly generated monocytes are dynamically remodeled during SIV infection and provide new insight into the mechanisms linking dysregulated monopoiesis to CNS disease.

These findings extend our previous work showing that expansion of circulating BrdU⁺ monocytes is a sensitive predictor of disease progression in the SIV macaque model. We previously demonstrated that increased percentages and absolute numbers of BrdU⁺ monocytes correlate with a shorter time to AIDS, more rapid disease progression, greater severity of SIVE neuropathology, and increased viral burden and inflammation within the CNS (Burdo et al., 2010; Hasegawa et al., 2009). The present study builds upon those observations by identifying the transcriptional programs that distinguish these pathogenic monocytes and may underlie their association with neuroinvasion and CNS injury.

Monocytes are traditionally divided into classical, intermediate, and nonclassical subsets based on expression based on expression of CD14 and CD16. In untreated HIV infection, the proportion of CD16+ monocytes increased in patients with severe HAND (Pulliam et al., 1997). Subsequent efforts have focused on transcriptomic changes within classical, intermediate, and nonclassical monocytes. But recent hematopoiesis studies have demonstrated that CD14 and also CD16 expression is insufficient to capture monocyte heterogeneity, where four transcriptionally distinct monocyte subtypes can be identified in steady-state conditions (Villani et al., 2017) and additional monocyte subsets are produced in response to inflammation (Swann et al., 2024). In this study, we sought to define the identity of BrdU+ monocytes that predict severe CNS histopathology in a primate model of HIV infection.

Differential expression profiles of SIVE and SIVE-predictive BrdU+ monocytes are consistent with an immature monocyte identity. Terminally, SIVE BrdU+ monocytes downregulated GO processes associated with inflammation, cell activation, and extravasation. SIVE-predictive BrdU+ monocytes downregulated interferon signaling, antigen presentation, and NF-kB signaling pathways relative to BrdU+ monocytes in SIVE non-progressors. Others have reported that the mobilization of immature monocytes correlates with increased inflammation by CD8 T-cell activation, increased viral load, and lower CD4 T-cell counts (Sui et al., 2017). Our data are consistent with an increase in %BrdU+ monocytes in SIVE progressors that may be driven by increased production of such immature monocytes, which appear to have an immune suppressive phenotype. This is also consistent with a higher percentage of BrdU+ monocytes in the blood at 24 h post i.v. BrdU inoculation.

Myeloid-derived suppressor cells (MDSCs) are a subtype of immature monocytes characterized by immunosuppressive function and decreased expression of HLA-DR (Bergenfelz and Leandersson, 2020). Transcriptionally, SIVE BrdU+ monocytes have upregulated IDO1, an enzyme that suppresses immune responses and inhibits pro-inflammatory monocyte maturation by scavenging reactive oxygen species(Ju et al., 2021; Okła et al., 2019; Tattevin et al., 2010). SIVE BrdU+ monocytes also have upregulated CD13, which is also associated with MDSC identity (Zhang et al., 2013, 2017). The downregulation of CD56 that we find may be explained by the expansion of immunosuppressive monocytes, which do not express CD56, relative to rare populations of CD56+ monocytes (Campana et al., 2022; Sconocchia et al., 2005). The observed upregulation of the immunosuppressive surface marker LILRB4 in SIVE-predictive BrdU+ monocytes is consistent with the idea that SIVE-predictive and SIVE BrdU+ monocytes are enriched for MDSCs (Gong et al., 2024; Lu et al., 2009). SIVE BrdU+ monocytes expressed lower levels of cell-surface HLA-DR, consistent with a potential MDSC identity, although HLA-DR expression did not statistically differentiate SIVE from SIVnoE BrdU+ monocytes. Bissel et al. (2006) has previously reported decreased HLA-DR expression on monocytes prior to the development SIVE (Bissel et al., 2006).

If SIVE and SIVE-predictive BrdU+ monocytes are MDSCs, this identity may explain why the histopathology of severe HAND – a condition associated with chronic inflammation – is characterized by an increase of CD163+CD206+ “anti-inflammatory” macrophages in the brain (Burdo et al., 2015). MDSCs preferentially differentiate into CD163+ macrophages *in vitro* (Bayik et al., 2018), correlate with the number of CD163+ macrophages in human cancer studies (Lv et al., 2022), and promote the polarization of macrophages towards a M2 phenotype in mouse (Sinha et al., 2007), which are identified by CD163 expression in primates (Ashley et al., 2016). In support of an MDSC identity, we have previously shown that at 21 DPI CNS BrdU+ macrophages are overwhelmingly MAC387 (Soulas et al., 2011), and others have found that expression of the S100A8 and S100A9 subunits that make up MAC387 is 20-fold greater in MDSCs vs non-MDSC monocytes (Zhao et al., 2012). Further phenotyping, functional assays, and single-cell RNA-seq are necessary to test the hypothesis that the SIVE and SIVE-predictive BrdU+ monocytes are enriched for MDSCs.

We have previously shown that a 2-to-10-fold increase in the %BrdU+ monocytes at 21DPI is a reliable and sensitive predictor of rapid AIDS and development of SIVE (Burdo et al., 2010; Hasegawa et al., 2009). Increased monocytopoietic output requires hematopoietic stem and progenitor cells (HSPCs) to undergo repeated cell replication, increased transcription, and increased metabolic output. Such demands can cause DNA damage, which skews HPSCs towards myeloid lineages at the expense of lymphoid lineages (Wang et al., 2023). In this study, we found that BrdU+ SIVE monocytes have a greater burden of structural variants compared to their SIVnoE counterparts. Interestingly, we used a spermidine analog mitoguazone or MGBG (methylglyoxal bis[guanylhydrazone]), and found that despite high plasma virus, it blocked the development of AIDS (unpublished data) and decreases the severity and prevalence of cardiovascular pathology and peripheral neuropathy in CD8 lymphocyte-depleted SIV infected monkeys (Lakritz et al., 2017). It is interesting to note in these studies the MGBG is only taken up by macrophages and not lymphocytes (Jin et al., 2015; Walker et al., 2017). Eight of the top 10 genes with structural variants were exclusive to BrdU+ SIVE monocytes and encoded extremely long genes - consistent with random DNA damage resulting from repeated replication of HSPCs. The upregulation of the DNA damage repair enzyme PARP2 in BrdU+ of SIVE progressors vs. non-progressors supports the notion that sustained monopoiesis results in mutations in SIVE-predictive BrdU+ monocyte progenitors and progeny. DNA damage caused by infection of monocytes is unlikely to be responsible for this DNA damage response, as HIV infection of monocytes is rare (Massanella et al., 2019).

In this study, we sought to transcriptionally characterize BrdU+ monocytes that predict the development of AIDS and the prevalence of SIVE. We provide evidence that SIVE-predictive and SIVE BrdU+ monocytes are an immature monocyte subpopulation that may be immunosuppressive. Increased mutational burden in SIVE BrdU+ monocytes raise the possibility that BrdU+ monocyte production associated with SIVE can result in dysregulated monopoiesis and that such dysregulation occurs when macrophages of innate immunity become dysregulated and activated with loss of acquired immunity (CD4 loss and CD8 exhaustion) with HIV and SIV infection. Overall, this work supports the notion that BrdU+ monocytes in SIVE progressors and terminal SIVE animals are enriched for a distinct monocyte subpopulation that is less frequent or absent in SIVnoE and non-infected animals. Future efforts to study HAND in relation to monocytes and macrophages should move beyond traditional CD14 and CD16-based classification of monocytes and towards finer-grained monocyte subtyping.

## Methods

### Animals

Male Indian-origin rhesus macaques (RMs; *Macaca mulatta*), aged 4–14 years, were used in this study (**Table S1**). All animals were confirmed free of SIV, HIV-2, STLV-1, and SRV at baseline. All procedures were approved by the Tulane University Institutional Animal Care and Use Committee and conducted in accordance with the NIH *Guide for the Care and Use of Laboratory Animals* and the Weatherall report. Routine procedures, including physical exams and blood draws, were performed under ketamine anesthesia to minimize discomfort. Animals were infected i.v. with SIVmac251 and underwent CD8^+^ cell depletion using the rhesus chimeric anti-CD8 antibody M-T807 (10 mg/kg subcutaneously on day 6, and 5 mg/kg intravenously on days 8 and 12 post-infection) to promote rapid AIDS progression and consistent development of SIVE (in >85% of persistently CD8-depleted animals). ART, consisting of oral raltegravir (22 mg/kg), subcutaneous tenofovir (20 mg/kg), and emtricitabine (30 mg/kg), was initiated at 21 days post-infection (DPI) and continued daily until euthanasia at either 12 or 14 weeks of treatment.

The thymidine analog 5-bromo-2′-deoxyuridine (BrdU; Sigma-Aldrich, St. Louis, MO) was used to label cells undergoing DNA synthesis. Animals received intravenous injection of BrdU (60 mg/kg) at 20 DPI and again 24 h prior to necropsy. Blood samples were collected 24 h post BrdU administration to assess monocyte turnover measured by incorporation of BrdU in monocyte precursors in BM that emigrate to blood at that timepoint (Burdo et al., 2013a). These data are consistent with the monoblast and the promonocyte that can divide in the BM but not blood or parenchymal tissues (Volkman, 1976). Some animals had additional BrdU timepoints. Animals were humanely euthanized upon developing AIDS-related criteria, including significant weight loss (>15% in 2 weeks or >30% in 2 months), persistent anorexia, unresponsive opportunistic infections, severe diarrhea, progressive neurological signs, or notable cardiac/pulmonary symptoms. SIVE diagnosis was confirmed by the presence of SIV RNA-positive cells, macrophage infiltration, and multinucleated giant cells (MNGCs) in the brain parenchyma.

### PBMC Collection and Fluorescence-Activated Cell Sorting

At 21 DPI and necropsy, EDTA-anticoagulated blood was collected, and peripheral blood mononuclear cells (PBMCs) were isolated via Ficoll-Paque density gradient centrifugation (GE Healthcare) and cryopreserved at −150°C in 10% DMSO 90% FBS. For fluorescence-activated cell sorting (FACS), PBMCs were thawed in two changes of 20% FBS-BSA, resuspended in 100 μL of wash buffer (1% BSA in PBS + 1:100 RNasin Plus, Promega Cat# N2611), then stained with HLA-DR PE-Texas Red (6:100), CD16-PECY7 (0.6:100), CD3 APC-CY7 (15:100), CD20 APC-CY7 (3:100), CD14 Pacific Blue (7.5:100), and LIVE/DEAD Fixable Aqua Dead Cell Stain (3:100, Invitrogen). Cells were incubated with antibody for 15 min at room temperature with rocking. Cells were washed with 1 mL of wash buffer, then fixed and permeabilized by adding 500 μL of BD Cytofix/Cytoperm and incubating for 20 min on ice. For intracellular BrdU staining, cells were washed with permeabilization buffer (0.1% saponin in wash buffer), resuspended in 100 μL of DNase (BD) diluted 1:10 in permeabilization buffer, incubated for one h at 37°C, washed with permeabilization buffer, and incubated with BrdU-FITC (BD Biosciences) for 20 min at room temperature. Cells were washed at the final time in wash buffer, then resuspended in a volume of 300 μL of wash buffer for sorting on a BD FACSAria IIIu (BD Biosciences). Single-stain controls for flow cytometry compensation analysis, heat-shocked cells as positive controls for dead cells (that are subsequently gated out in flow cytometry sorting), and fluorescence-minus-one controls for HLA-DR, Live/Dead, and BrdU were prepared for each run. To identify BrdU-labeled monocytes, we gated on the whole monocyte population by forward and side scatter, excluded doublets by FSC-A and FSC-H, excluded dead cells using a Live/Dead dye, selected for HLA-DR-positive cells, identified monocytes by CD14 and CD16 expression, then gated monocytes based on BrdU. Cells were sorted into BSA-coated tubes, centrifuged and aspirated, snap frozen in ethanol and dry ice, then stored at -80°C prior to RNA isolation and library preparation.

### RNA Isolation, Library Preparation, and Sequencing

RNA was extracted using the miRNeasy FFPE kit (Qiagen), modifying the proteinase K step to 56°C for 1 h to preserve RNA integrity. RNA quality was assessed by DV200 score using an Agilent Bioanalyzer 2100 (Agilent). Libraries were prepared using the SMARTer Stranded Total RNA-Seq v3 - Pico Input Mammalian (Takara) according to manufacturer instructions. Sequencing library quality was assessed using the Agilent Bioanalyzer 2100 (Agilent) and quantified using the Qubit 1X HS dsDNA High Sensitivity kit (Invitrogen). Sequencing was performed by the University of Florida Interdisciplinary Center for Biotechnology Research NextGen DNA sequencing core to a depth of approximately 300 million paired-end reads per sample on a NovaSeq X to achieve approximately 50 million exon-mapping reads per sample. To mitigate potential batch effects, BrdU+ and BrdU-infected and non-infected samples were included in each batch of FACS, RNA extraction, and library preparation.

### Data Processing

Raw reads were processed to remove low-quality sequences and adapters, and the resulting clean reads were aligned to the rhesus macaque reference genome (*NCBI Mmul_10*) using the STAR aligner. Gene annotation was conducted using resources provided by the Cebola Lab. Genes with fewer than four normalized counts across all samples were excluded from downstream analysis. Data normalization was performed using the precision weights approach called voom, which transforms count data to log2-counts per million (log_2_CPM) while estimating the mean–variance relationship to assign precision weights(Law et al., 2014). This method enables robust application of linear modeling frameworks via the *limma* package, effectively addressing heteroscedasticity in RNA-seq data. Limma voom was selected for its strength in handling complex experimental designs and moderate sample sizes, offering improved statistical power for identifying differentially expressed genes. Additionally, it supports advanced downstream analyses, including variance component score testing for gene set enrichment in longitudinal RNA-seq studies, as demonstrated by Agniel and Hejblum (2017) (Agniel and Hejblum, 2017). Differential expression analysis was performed using the limma pipeline in R (version 4.4.0), following normalization with the voom method, including variancePartition, edgeR, and doParallel packages. Voom-transformed expression data (log_2_CPM) were used to perform PCA to assess sample clustering. Gene expression heatmaps were generated using pheatmap after Z-score normalization. To visualize DEGs, volcano plots were generated using the EnhancedVolcano package, while heatmaps were constructed with ggplot2, pheatmap, and viridis. Venn diagrams were used to identify overlapping DEGs across different comparison groups.

### RNA-seq Enrichment Analyses

Enrichment analyses were performed to examine biological processes, molecular functions, cellular components, and disease ontology using Metascape and the ClueGO plug-in (version 2.5.8) (Bindea et al., 2009; Zhou et al., 2019). Significance was determined using adjusted p-values < 0.05, with multiple comparison corrections applied via the Benjamini-Hochberg method. For ClueGO analysis, signaling pathways were sourced from KEGG, Reactome, and WikiPathways databases, and enrichment was assessed using a two-sided hypergeometric test with Bonferroni step-down correction. Term associations were established based on a κ score threshold of 0.3. ClueGO integrates Gene Ontology (GO) terms and pathway data from KEGG and BioCarta. Key DEGs were identified through network topology analyses, including betweenness, closeness, and degree centrality, using the CytoHubba application in Cytoscape. Additionally, transcription factors (TFs) regulating gene expression changes were identified using ChEA3 (Keenan et al., 2019). DEGs were input into ChEA3, which assessed their association with known TF target datasets through Fisher’s Exact Test, using a reference background size of 20,000 genes.

### Quantitative RT-PCR

Two genes were selected for validation due to their biological relevance to SIV pathogenesis, monocyte function, and neuroinflammation: *SPI1* (PU.1), a transcription factor essential for monocyte differentiation and implicated in HIV-associated neuroinflammation; and *PARP2*, associated with monocyte activation, oxidative stress, and neuroinflammatory responses observed in SIVE. Total RNA isolated from CD14⁺ monocytes was used to validate gene expression by quantitative reverse transcription PCR (qRT-PCR). Reactions were performed in singlet for each sample using the Applied Biosystems TaqMan™ RNA-to-CT™ 1-Step Kit and TaqMan™ Gene Expression Assays for SPI1 (Rh02844466_m1), PARP2 (Rh01003788_m1), and the endogenous control β-actin (Hs00357333_g1) (Thermo Fisher Scientific). Thermal cycling conditions were as follows: 48°C for 15 min, 95°C for 10 min, followed by 40 cycles of 95°C for 15 s and 60°C for 1 min on a QuantStudio 7 Pro PCR system. For analysis, cycle threshold (Ct) values of target genes were normalized to β-actin to generate ΔCt values. Relative gene expression was determined by calculating ΔΔCt values relative to samples from SIV-negative animals, and fold changes were calculated using the 2^−ΔΔCt^ method.

### Structural variant calling

To identify and compare structural variants (SVs) in BrdU+ and BrdU-CD14⁺ monocytes isolated from brain tissues collected at necropsy from three AIDS-SIVE RMs, we employed a combined computational approach using Manta (v1.6.0), FreeBayes (v1.3.10), and DeepVariant (v1.5.0). High-quality paired-end sequencing data were aligned to the Macaca mulatta reference genome (*Mmul_10*) using BWA-MEM. Manta was used to detect large (>50 bp) structural variants, including deletions, insertions, duplications, and inversions, with default parameters optimized for germline analysis (Chen et al., 2016). To complement Manta’s calls and improve variant detection sensitivity and precision, FreeBayes and DeepVariant were applied to identify small indels (1-50 bp) and refine breakpoint resolution. Variants identified by these tools were filtered for high-confidence calls based on quality scores, read support, and allele frequency.

High-confidence variants from Manta, FreeBayes, and DeepVariant were filtered based on the following criteria: (a) phred-scaled quality score (QUAL > 30), minimum read support (≥ 10 supporting reads), (c) allele frequency thresholds to distinguish somatic vs. germline events, and (d) removal of low-complexity and repetitive regions using RepeatMasker annotations.

High-confidence variants from these tools were filtered to retain only those with a PASS flag in the FILTER column using bcftools (v1.13). Remaining variants were annotated with SnpEff (v5.0) against the *Macaca mulatta* Mmul_10.99 reference genome to predict functional effects (e.g., impact, gene name, biotype). Non-coding and artefactual annotations (e.g., snoRNAs, snRNAs, Y RNAs, 7SK, 7SL) were removed. We performed pairwise comparisons of SV profiles between BrdU⁺ and BrdU⁻ monocytes within each individual. Differentially mutated genes were identified using Fisher’s exact test, followed by FDR correction using the Benjamini-Hochberg method. Significant genes (q-value < 0.05) were visualized via volcano plots and bar plots, highlighting those enriched in BrdU⁺ cells, which may reflect proliferation- or activation-related genomic instability. VCF outputs were parsed using custom bash and R scripts to generate mutation matrices by gene. Normalized mutation counts were plotted using ggplot2, and top candidates were annotated and prioritized for downstream pathway enrichment analysis. Key mutated genes were identified based on network topology metrics—betweenness, closeness, and degree centrality—using the CytoHubba plugin in Cytoscape.

## Acknowledgment

The anti-CD8alpha [MT807R1] antibody used in this study was provided by the NIH Nonhuman Primate Reagent Resource (P40 OD028116 and U24 AI126683).

## Author contribution

HDN: Methodology, Validation, Formal Analysis, Data Curation, Writing – Original Draft, Visualization

AKD: Validation, Formal Analysis, Investigation, Data Curation, Writing – Original Draft, Visualization

CES: Investigation, Data Curation PA: Investigation

MNC: Investigation

MS: Methodology, Investigation

KCW: Conceptualization, Methodology, Resources, Data Curation, Writing – Review & Editing, Supervision, Project Administration, Funding Acquisition

W-KK: Methodology, Formal Analysis, Resources, Data Curation, Writing – Review & Editing, Supervision, Project Administration, Funding Acquisition

## Study Funding

This study was supported by funding from the National Institutes of Health, R01NS040237 (KCW), R01NS126091 (KCW & W-KK), and R01MH118139 (W-KK). The animal experiments were done at the TNBRC and were in part supported in part by the TNBRC P51 Base Grant (P51OD011104). MS and MNC were supported by the Stephany W. Holloway University Chair in AIDS Research.

## Data availability statement

Data and code will be made available freely from the corresponding authors upon reasonable request.

## Supplementary Figure

**Figure S1.**
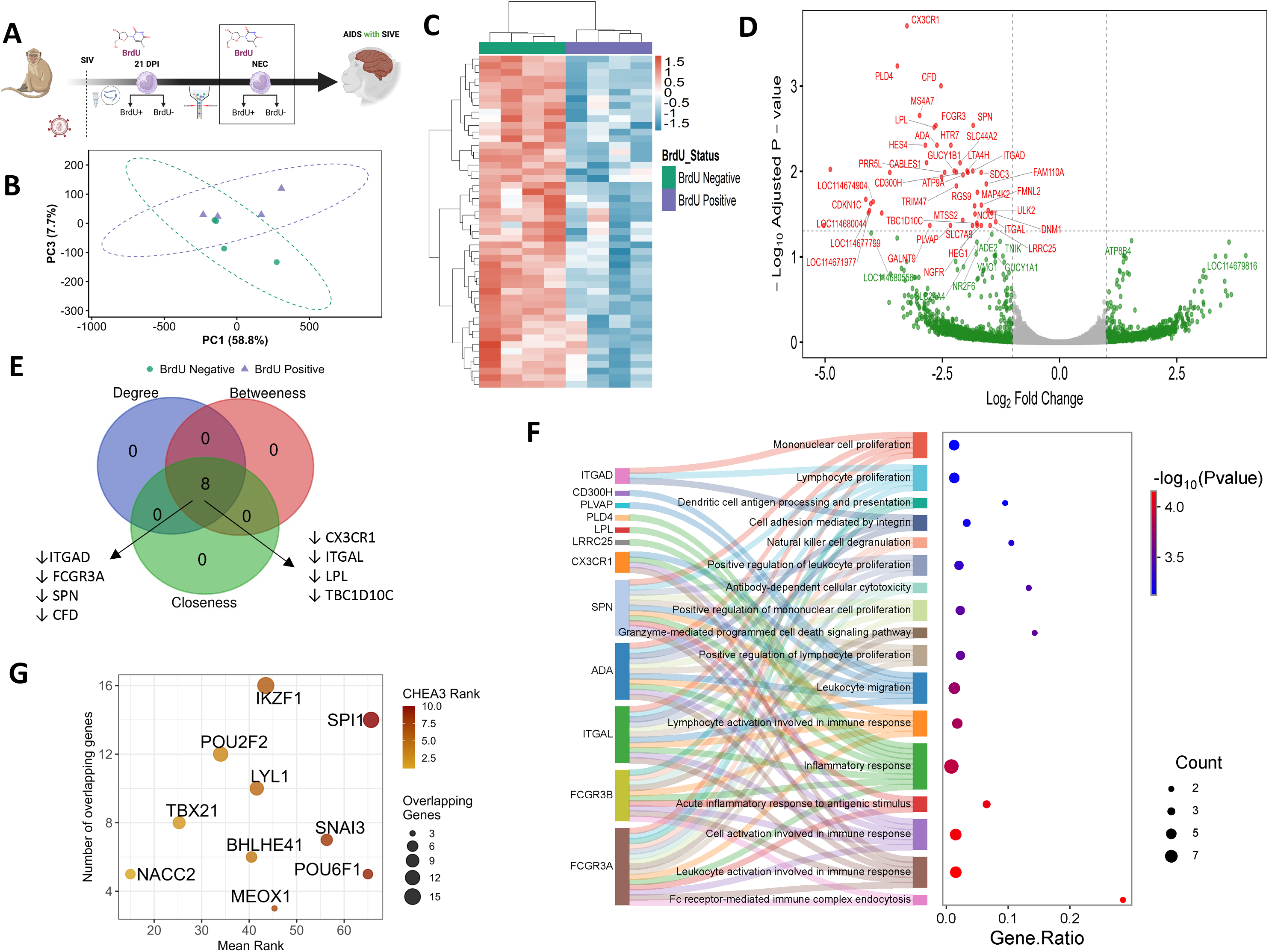
Transcriptomic profiling of BrdU⁺ and BrdU⁻ monocytes in SIV-associated encephalitis at necropsy. (**A**) Experimental design: SIV-infected rhesus macaques with encephalitis (SIVE) were administered BrdU before necropsy. BrdU⁺ and BrdU⁻ monocytes were sorted for RNA-seq analysis. (**B**) Principal component analysis (PCA) of BrdU⁺ and BrdU⁻ monocytes showing separation between groups (PC1 = 58.8%, PC3 = 7.7%). (**C**) Heatmap of the top 50 differentially expressed genes (DEGs) across samples. (**D**) Volcano plot of DEGs between BrdU⁺ and BrdU⁻ monocytes. Dashed lines indicate thresholds of |log2FC| = 1 and adjusted *P* = 0.05. Non-significant genes are shown in grey; genes with |log2FC| > 1 only in green; adjusted *P* < 0.05 only in blue; and significant genes meeting both criteria in red. (**E**) Network centrality analysis identifying hub genes associated with SIV pathogenesis based on degree, betweenness, and closeness metrics. (**F**) Sankey and dot plots showing Gene Ontology (GO) enrichment of DEGs. (**G**) Transcription factor (TF) enrichment analysis identifying key regulators using ChEA3 (ChIP-X Enrichment Analysis 3), ranked by overlap and mean rank; lower mean rank indicates higher regulatory significance.

**Figure S2.**
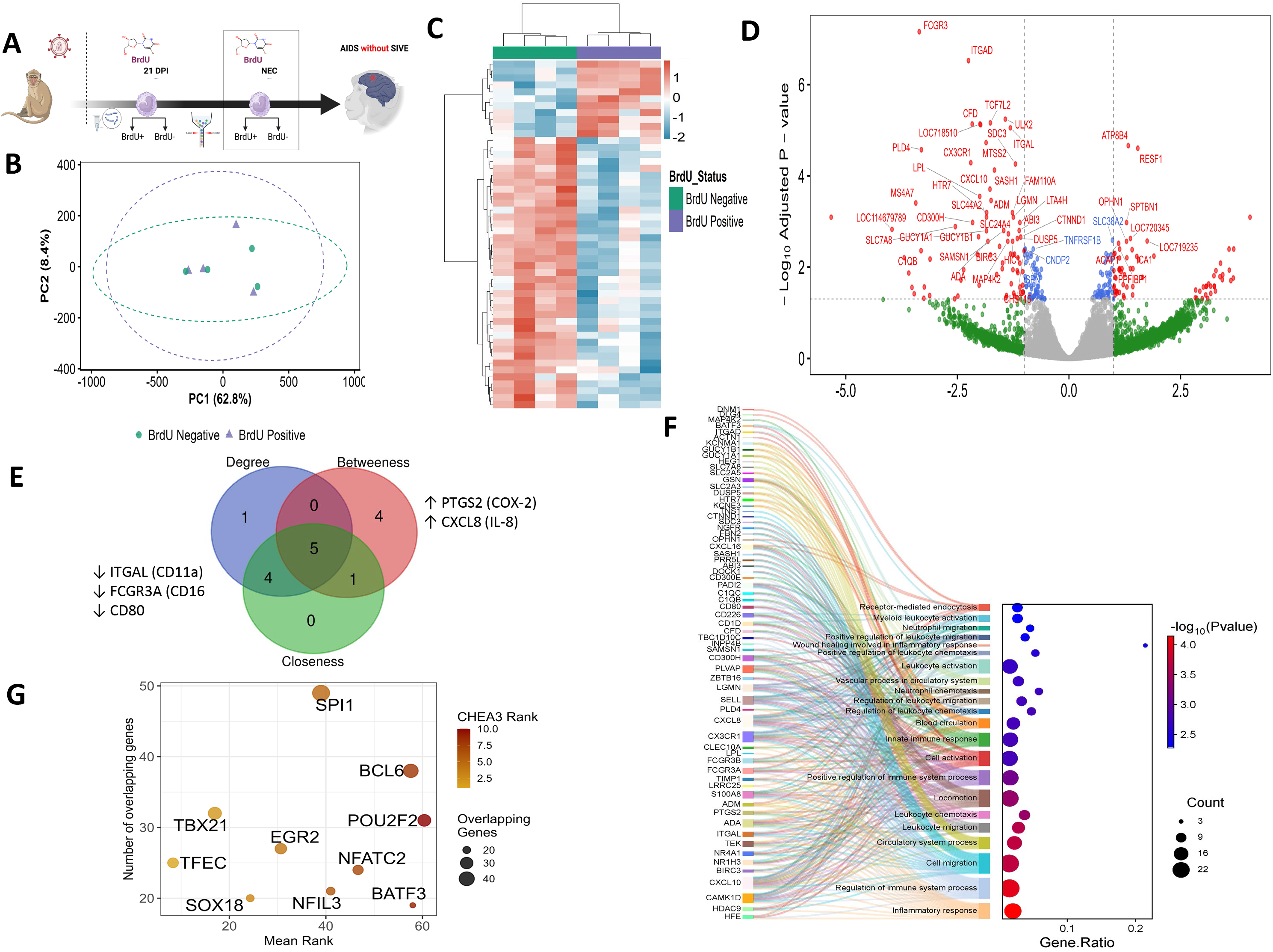
Transcriptomic profiling of BrdU⁺ and BrdU⁻ monocytes in SIV infection without encephalitis at the necropsy. (**A**) Experimental design: SIV-infected rhesus macaques without encephalitis (SIVnoE) were administered BrdU prior to necropsy. BrdU⁺ (proliferating) and BrdU⁻ (non-proliferating) monocytes were sorted for RNA-seq analysis. (**B**) Principal component analysis (PCA) of BrdU⁺ and BrdU⁻ monocytes showing separation between groups (PC1 = 62.8%, PC3 = 8.4%). (**C**) Heatmap of the top 50 differentially expressed genes (DEGs) across samples. (**D**) Volcano plot of DEGs between BrdU⁺ and BrdU⁻ monocytes. Dashed lines indicate thresholds of |log2FC| = 1 and adjusted *P* = 0.05. Non-significant genes are shown in grey; genes with |log2FC| > 1 only in green; adjusted *P* < 0.05 only in blue; and significant genes meeting both criteria in red. (**E**) Network centrality analysis identifying hub genes associated with SIV pathogenesis based on degree, betweenness, and closeness metrics. (**F**) Sankey and dot plots showing Gene Ontology (GO) enrichment of DEGs. (**G**) Transcription factor (TF) enrichment analysis identifying key regulators using ChEA3 (ChIP-X Enrichment Analysis 3), ranked by overlap and mean rank; lower mean rank indicates higher regulatory significance.

**Figure S3.**
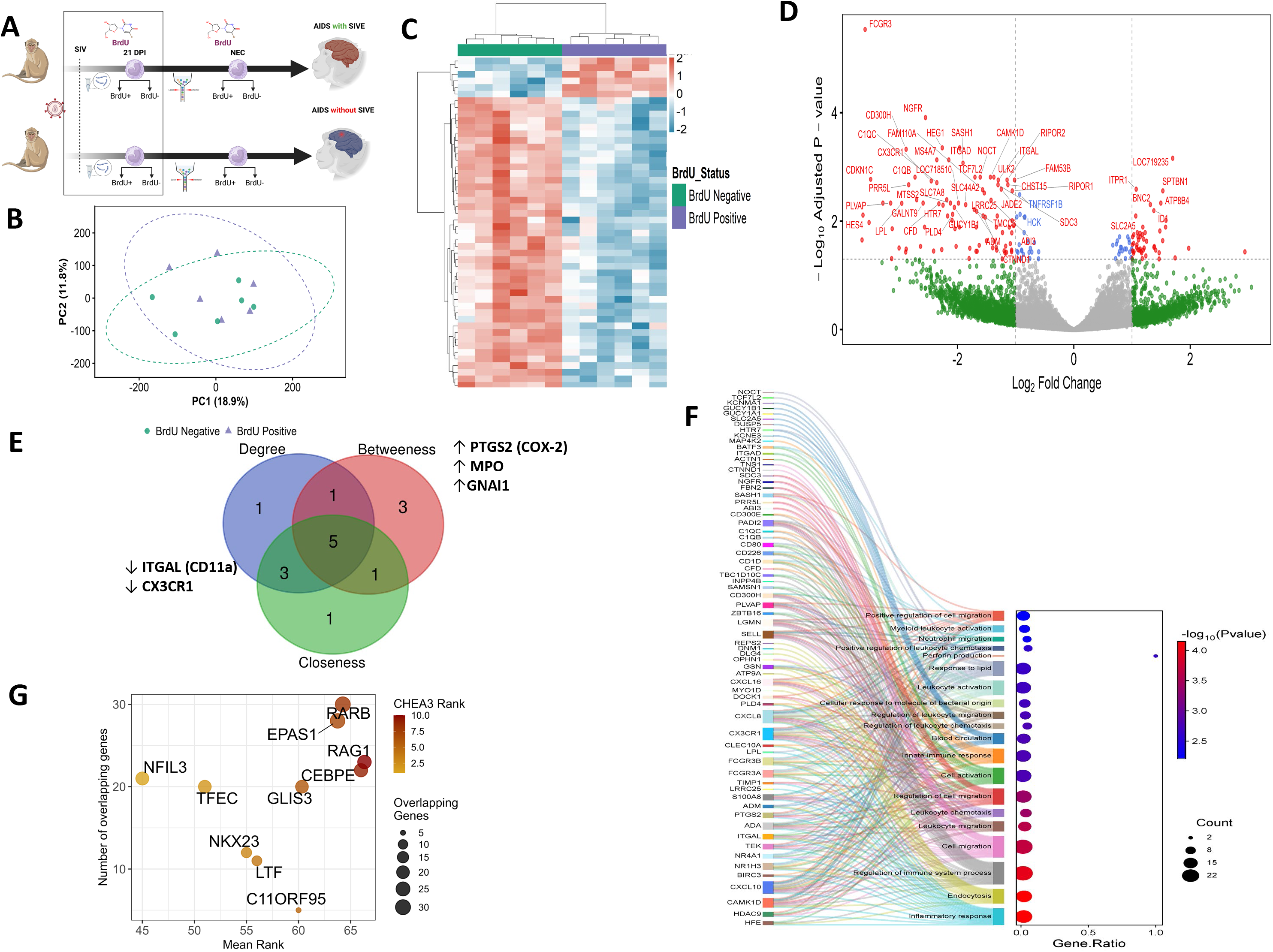
Transcriptomic profiling of BrdU⁺ and BrdU⁻ monocytes during acute SIV infection. (**A**) Experimental design: SIV-infected rhesus macaques during acute infection were administered BrdU at 21-day post infection (DPI). BrdU⁺ (proliferating) and BrdU⁻ (non-proliferating) monocytes were sorted for RNA-seq analysis. (**B**) Principal component analysis (PCA) of BrdU⁺ and BrdU⁻ monocytes showing separation between groups (PC1 = 18.9%, PC2 = 11.8%). (**C**) Heatmap of the top 50 differentially expressed genes (DEGs) across samples. (**D**) Volcano plot of DEGs between BrdU⁺ and BrdU⁻ monocytes. Dashed lines indicate thresholds of |log2FC| = 1 and adjusted *P* = 0.05. Non-significant genes are shown in grey; genes with |log2FC| > 1 only in green; adjusted *P* < 0.05 only in blue; and significant genes meeting both criteria in red. (**E**) Network centrality analysis identifying hub genes associated with SIV pathogenesis based on degree, betweenness, and closeness metrics. (**F**) Sankey and dot plots showing Gene Ontology (GO) enrichment of DEGs. (**G**) Transcription factor (TF) enrichment analysis identifying key regulators using ChEA3 (ChIP-X Enrichment Analysis 3), ranked by overlap and mean rank; lower mean rank indicates higher regulatory significance.

**Figure S4.**
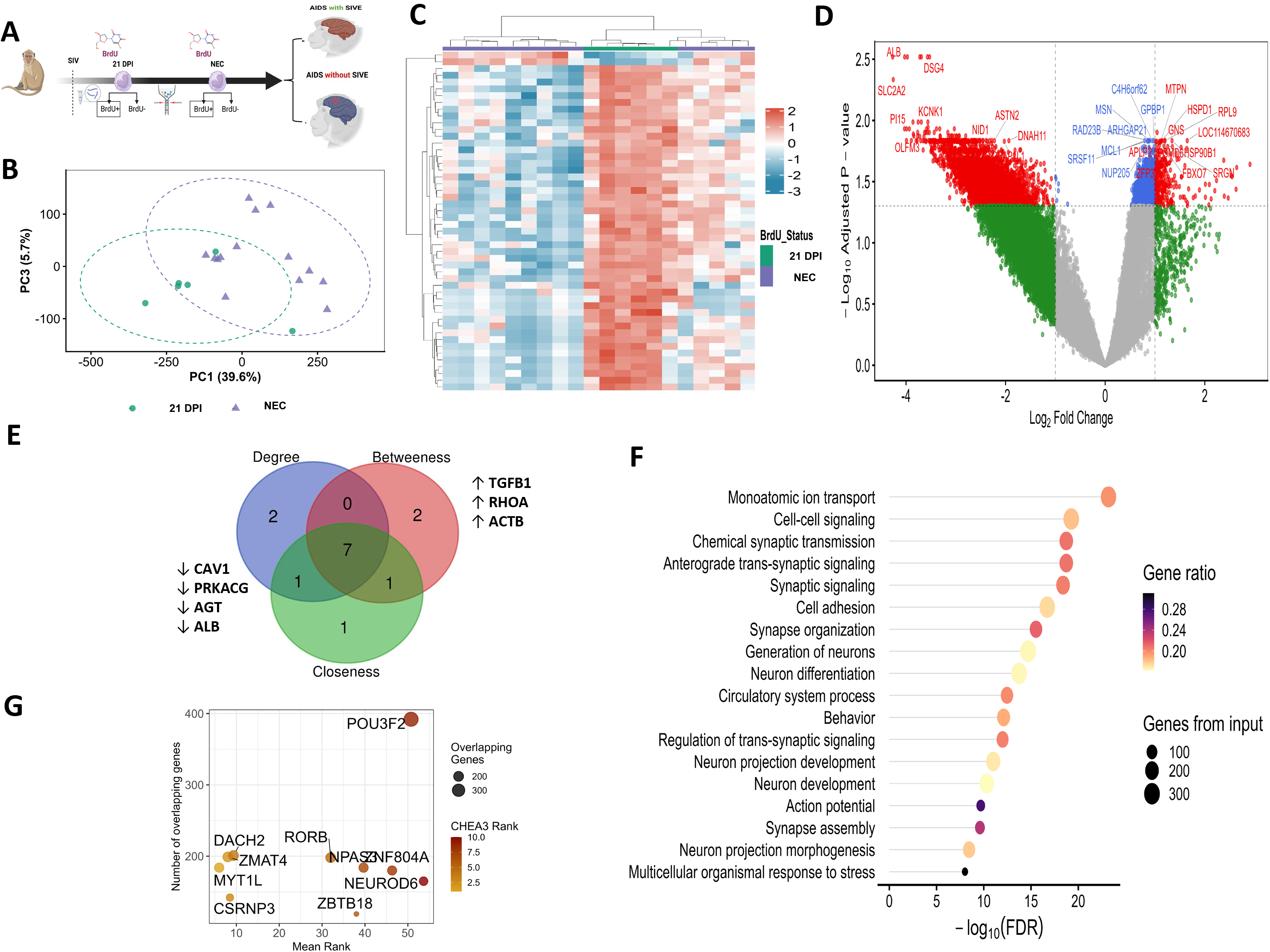
Transcriptional reprogramming of proliferating monocytes between early and terminal SIV infection. (**A**) Experimental design: SIV-infected rhesus macaques during acute infection (at 21-day post infection (DPI)) and necropsy were administered BrdU. BrdU⁺ (proliferating) and BrdU⁻ (non-proliferating) monocytes were sorted for RNA-seq analysis. BrdU⁺ monocytes were used to downstream analysis. (**B**) Principal component analysis (PCA) of BrdU⁺ monocytes during 21 DPI and NEC showing separation between groups (PC1 = 39.6%, PC3 = 5.7%). (**C**) Heatmap of the top 50 differentially expressed genes (DEGs) across samples. (**D**) Volcano plot of DEGs between BrdU⁺ monocytes during 21 DPI and NEC. Dashed lines indicate thresholds of |log2FC| = 1 and adjusted *P* = 0.05. Non-significant genes are shown in grey; genes with |log2FC| > 1 only in green; adjusted *P* < 0.05 only in blue; and significant genes meeting both criteria in red. (**E**) Network centrality analysis identifying hub genes associated with SIV pathogenesis based on degree, betweenness, and closeness metrics. (**F**) Sankey and dot plots showing Gene Ontology (GO) enrichment of DEGs. (**G**) Transcription factor (TF) enrichment analysis identifying key regulators using ChEA3 (ChIP-X Enrichment Analysis 3), ranked by overlap and mean rank; lower mean rank indicates higher regulatory significance.

**Figure S5.**
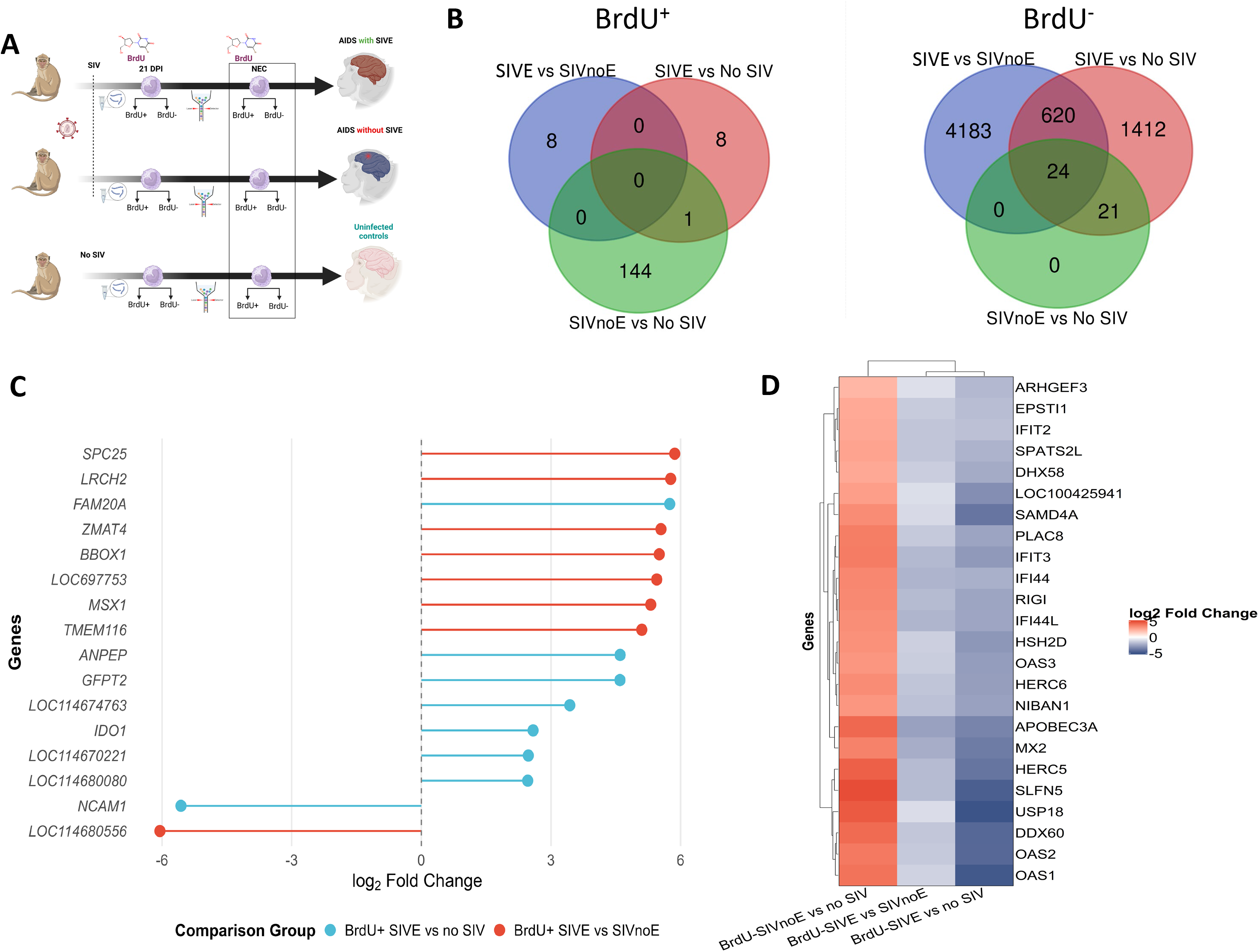
Distinct and Shared Transcriptional Signatures in BrdU⁺ and BrdU⁻ Monocytes during SIV Infection. **(A)** Experimental design: BrdU⁺/ BrdU^-^ monocytes from SIVE animals (n=3) were compared to BrdU⁺/ BrdU^-^ monocytes from SIVnoE animals (n=4) and uninfected (no SIV) controls at necropsy (n=6). **(B)** Venn diagram showing unique and shared genes differentially expressed in BrdU⁺ SIVE monocytes/ BrdU^-^ SIVE monocytes compared to SIVnoE and control groups. **(C)** Lollipop plot of the 16 unique genes with log2FC values (all adjusted P-values<0.05). **(D)** Heatmap of 24 shared genes in BrdU^-^ monocytes at necropsy across conditions. Data represent log2FC (all adjusted P-values<0.05).

**Fig S6.**
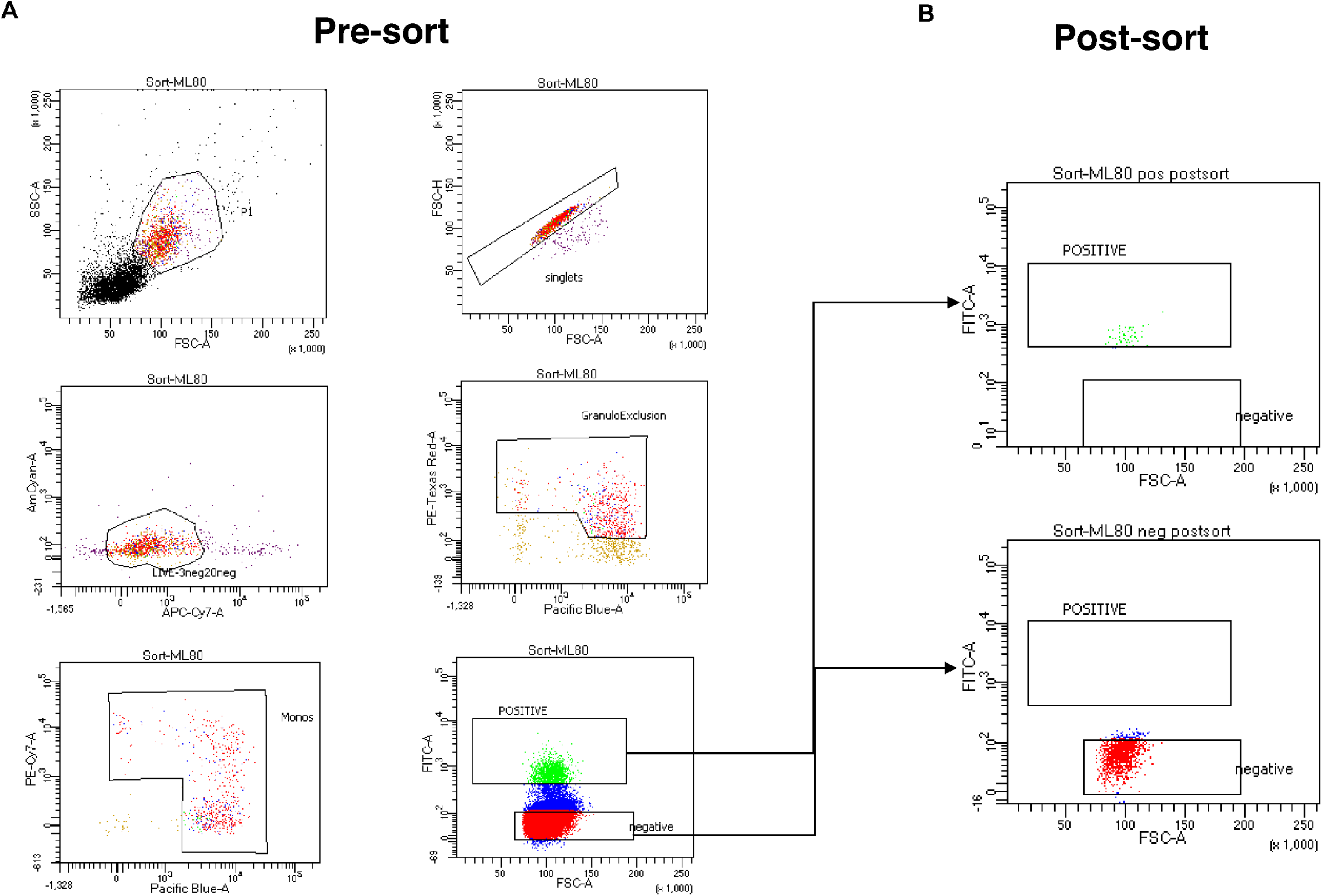
BrdU+ and BrdU-monocytes were sorted with extremely high purity. A) Raw data from flow cytometer demonstrating sort gating for BrdU+ monocytes B) Post-sort purity check for the sample sorted in panel A, demonstrating purity exceeding 99.9% for the BrdU+ and BrdU-fractions.

**Figure S7.**
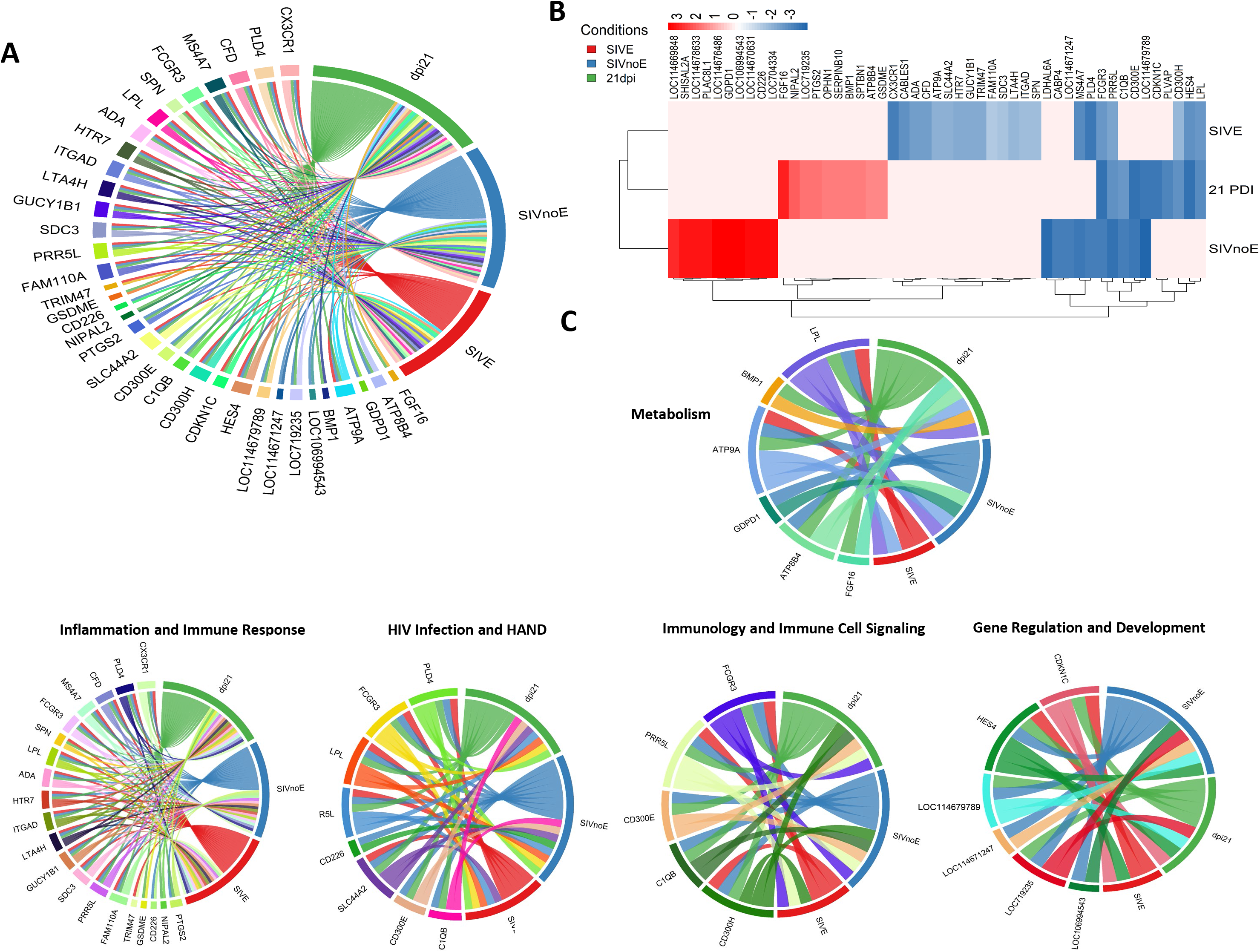
Shared and stage-specific transcriptional programs of proliferating monocytes across SIV infection. (**A**) Chord diagram of differentially expressed genes (DEGs) from three comparisons (SIVE necropsy, SIVnoE necropsy, and 21 DPI), highlighting a core set of genes common to all conditions. (**B**) Heatmap of shared and unique genes across early and late stages of SIV infection. (**C**) Functional enrichment analysis of condition-specific gene sets, showing involvement in metabolism, inflammation, immune response, HIV/HAND-related pathways, and gene regulation.

**Figure S8.**
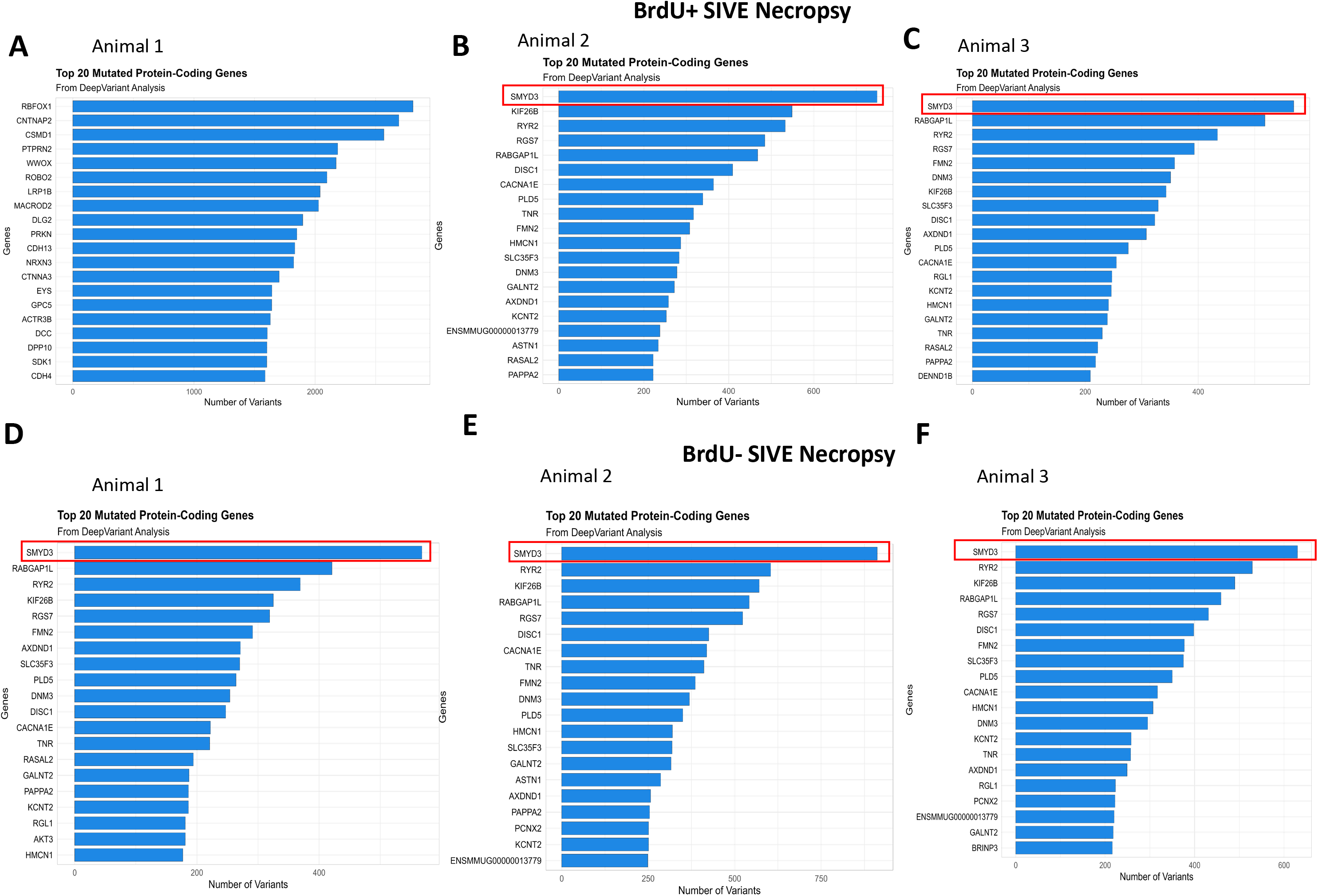
*SMYD3* structural variants in SIVE and SIVnoE BrdU⁺ monocytes. (**A-C**) Top 10 genes harboring SVs based on SnpEff annotation in SIVE BrdU⁺ monocytes. (**D-F**) Top 10 genes harboring SVs based on SnpEff annotation in SIVE BrdU^-^ monocytes.

## Supplementary Tables

**Table S1.** Animal information used in this study.

**Table S2.** Shared and stage-specific transcriptional programs of proliferating monocytes across SIV infection.

**Table S3.** Transcriptomic profiling of BrdU⁺ and BrdU⁻ monocytes in SIV-associated encephalitis at necropsy.

**Table S4.** Transcriptomic profiling of BrdU⁺ and BrdU⁻ monocytes in SIV infection without encephalitis at the necropsy.

**Table S5.** Transcriptomic profiling of BrdU⁺ and BrdU⁻ monocytes during early SIV infection.

**Table S6.** Transcriptional reprogramming of proliferating monocytes between early and terminal SIV infection.

**Table S7.** Early transcriptional signatures in BrdU⁺ monocytes predict progression to SIV-associated encephalitis.

**Table S8.** Distinct and Shared Transcriptional Signatures in BrdU⁺ and BrdU⁻ Monocytes during SIV Infection

**Table S9.** Genes show significantly different mutation burdens between BrdU⁺ and BrdU⁻ cells.

## Notes

### Competing Interest Statement

The authors have declared no competing interest.

